# Mechanical signals inhibit growth of a grafted tumor *in vivo*: Proof of Concept

**DOI:** 10.1101/045534

**Authors:** Rémy Brossel, Alexandre Yahi, Stéphane David, Laura Moreno Velasquez, Jean-Marc Guinebretière

## Abstract

In the past ten years, many studies have shown that malignant tissue has been “normalized” *in vitro* using mechanical signals. We apply the principles of physical oncology (or mechanobiology) *in vivo* to show the effect of a “constraint field” on tumor growth. The human breast cancer cell line, MDA MB 231, admixed with ferric nanoparticles was grafted subcutaneously in Nude mice. The magnetizable particles rapidly surrounded the growing tumor. Two permanent magnets located on either side of the tumor created a gradient of magnetic field. Magnetic energy is transformed into mechanical energy by the particles acting as “bioactuators”, applying a constraint field and, by consequence, biomechanical stress to the tumor. This biomechanical treatment was applied 2 hours/day during 21 days, from Day 18 to Day 39 following tumor implantation. The study lasted 74 days. Palpable tumor was measured two times a week. There was a significant *in vivo* difference between the median volume of treated tumors and untreated controls in the mice measured up to D 74 (D 59 + population): (529 [346; 966] mm^3^ vs 1334 [256; 2106] mm^3^; p=0.015), treated mice having smaller tumors. The difference was not statistically significant in the group of mice measured at least to D 59 (D 59 population). On *ex vivo* examination, the surface of the tumor mass, measured on histologic sections, was less in the treated group, G1, than in the control groups: G2 (nanoparticles, no magnetic field), G3 (magnetic field, no nanoparticles), G4 (no nanoparticles, no magnetic field) in the D 59 population (Median left surface was significantly lower in G1 (5.6 [3.0; 42.4] mm^2^, p=0.005) than in G2 (20.8 [4.9; 34.3]), G3 (16.5 [13.2; 23.2]) and G4 (14.8 [1.8; 55.5]); Median right surface was significantly lower in G1 (4.7 [1.9; 29.2] mm^2^, p=0.015) than in G2 (25.0 [5.2; 55.0]), G3 (18.0 [14.6; 35.2]) and G4 (12.5 [1.5; 51.8]). There was no statistically significant difference in the day 59+ population. This is the first demonstration of the effect of stress on tumor growth *in vivo* suggesting that biomechanical intervention may have a high translational potential as a therapy in locally advanced tumors like pancreatic cancer or primary hepatic carcinoma for which no effective therapy is currently available.

## INTRODUCTION

In the great majority of cases, a cancerous mass is a complex of tissues: an extracellular matrix (ECM), stroma, and the cancer cells derived from epithelium. In a clinical cancer, ECM is rigid whereas the tumor tissue is softer than the tissue of origin (1). The stiffness of a cell/tissue is determined by its “Young’s modulus” (E) which is obtained by measuring resistance to deformity of the cell/tissue in response to a force applied for a given surface area. E is expressed in Pascal (Pa). The measurement of stiffness with Young’s Modulus E was first performed *in vitro* on cultured cells. This has become a diagnostic tool for effusion cells (2) cell lines (3-5) or circulating cells (6). Depending on the technique and origin of the cells, the absolute values differ, but a common pattern can be seen with E greater than 1 kiloPascal (kPa) for normal cells with a relatively large spread of values. Following transformation to cancer, the same cells exhibit an E value of less than 1 kPa with less variation (7-9).

Recently *ex vivo* measures on breast biopsies maintained viable *ex vivo* have confirmed the *in vitro* findings (10); Normal breast tissue has a unimodal stiffness distribution of 1.13 +/-0.78 kPa. Cancer biopsies have a bimodal distribution with a first peak at 0.57+/-0.16 kPa representing the cancer tissue. A second peak at 1.99+/-0.73 kPa represents the ECM (9); The differential between cancer and non cancer is around 30-40 Pa (7-9). These studies also provided values for neoangiogenic tissue (10). *In vitro* and *ex vivo* measurements have in common the extensive use of Atomic Force Microscope and other approaches that have allowed us to have access to accurate measurement of cell and tissue mechanical parameters like Young’s module. This is useful to estimate the range of forces to be applied to a tumor to be able to overcome its rigidity. The delicate balance of forces between the tumor, the ECM, and the normal tissue surrounding the ECM is poorly understood. The best known of these mechanisms is the translation of mechanical signals to biological changes while the experimental study of the translation of mechanical cues to mechanical changes is scarce in the literature (11). The relationship in time and space of the forces controlling cancer and the surrounding normal tissues is yet to be studied. A tumor, therefore, is a complex ecosystem in which change from a non-malignant epithelial Euclidian architecture to a fractal architecture of cancer on a microscopic and macroscopic scale is accompanied by a change in mechanical/biochemical signals and a change in the physical properties of tissues in the cancerous complex (12-14). Cells and tissues, normal and cancerous, are continuously exposed to compression, surface / interface tension, hydrostatic and osmotic pressure, shear stress, etc… and more generally to constraint fields. In the last decade, the biophysical mechanisms associated with carcinogenic change and in ECM control of tissue growth and modeling (15) have become better understood (16-18). Three-dimensional (3D) culture systems have allowed the study of growth and apoptosis. It has been shown, for instance, that pressure applied to a tumor *in vitro* in different experimental setups delays tumor growth by changing cell death (apoptosis) and cell division in space and time (19-23). The first demonstration of the action of stress (surface tension) on the phenotype of a breast acinus was reported in 2005 (19). Others have shown the effect of osmotic pressure or compressive stress on a spheroid (20-22) or the effect of gravity on cells in a 2D cell culture (23). In those experiments, pressure was applied on tissue externally by modifying the environment that was then considered as being equivalent to the ECM. Tumor volume, rates of cell division and apoptosis are therefore measurable *in vitro*. They correlate with the effect of stress.

To apply a force to a malignant tumor *in vivo*, it was necessary to design an experimental device allowing pressure to be applied only to the tumor tissue using a process which delivered a constant pressure, locally. We present here the first *in vivo* experiment using pressure applied to a cancer mass of human origin to modify its growth^1^. This disruptive innovation consists of the capacity to modify tumor growth with a constraint field usable *in vivo*. The principle is illustrated on Fig. 1A showing a 2D culture *in vitro*.

**Fig 1:**
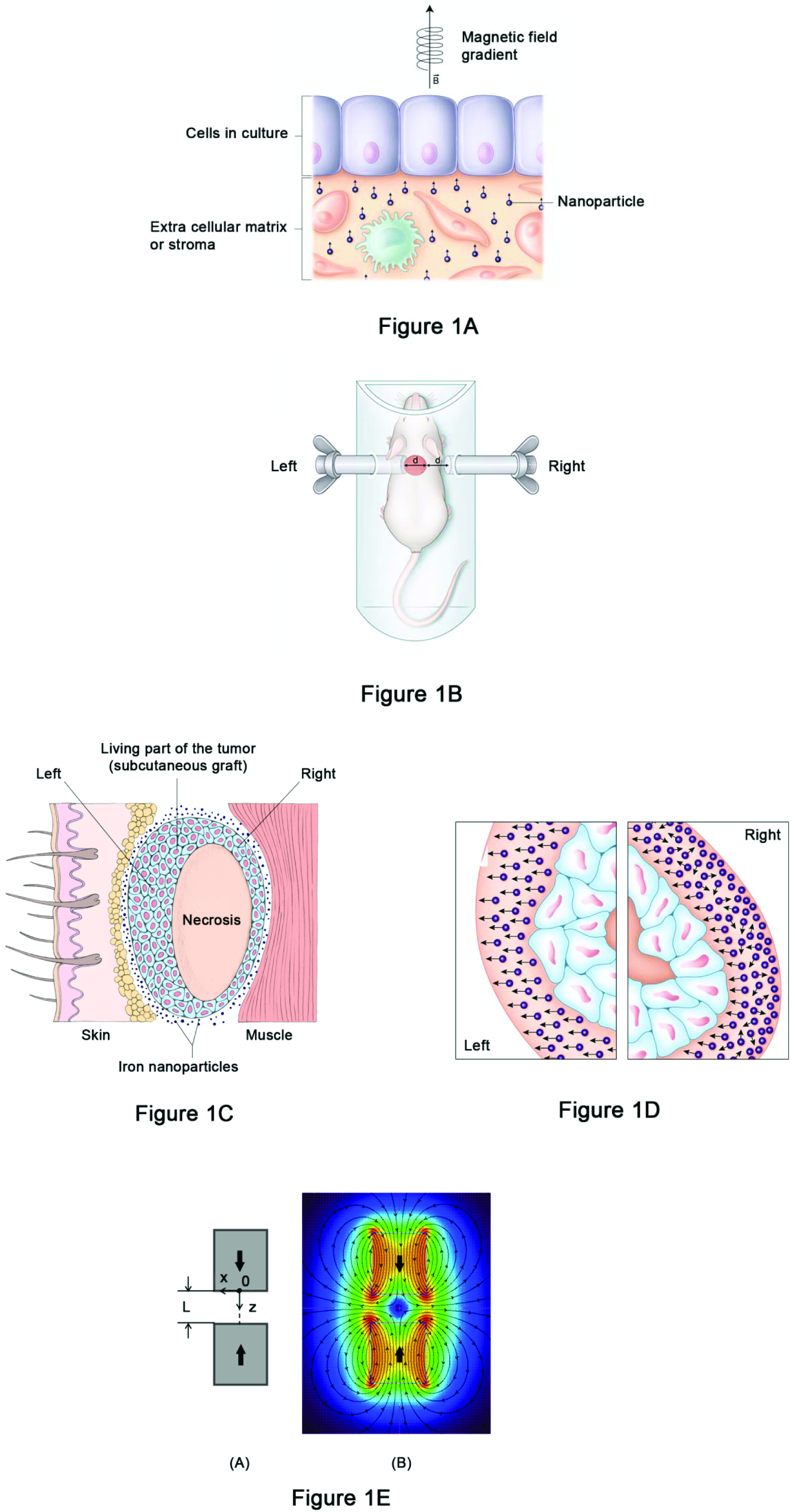
Fig. 1A: Principle of the use of a *in vivo* constraint field. Generation of a field of constraint on a 2D cell culture: each particle subjected to a magnetic field gradient generates a force vector. Altogether these vectors create a field of constraint pushing / pulling the cells depending on the orientation of the field. Fig. 1B: Proof of Concept. Schematic representation of the experimental setup with the animal in the airgap of two magnets (Magnets, tumor not at scale). Fig. 1C: Schematic representation of the living parts (right and left) and the necrosis of the subcutaneous grafted tumor. Fig. 1D: Close up of right and left parts. Force vectors are oriented towards the magnetic field or not depending on the distance to the magnet. Fig. 1E: Magnets in opposite configuration and typical gradient variation along the oz axis. (A) - configuration scheme; (B) - related indicative mapping of the magnetic field from *Vizimag* simulation.

To be able to apply the field *in vivo*, two permanent magnets located on either side of the tumor delivered a gradient of magnetic field. Magnetic energy is transformed into mechanical energy by the magnetized particles acting as “bioactuators”, applying a constraint field, and by consequence biomechanical stress to the tumor.

## STUDY RATIONALE

The action of a constraint field on a tissue is the result of biological consequences, only some of which are known: a very large number of biochemical signals are activated simultaneously either by conformational change as in changes in histone/DNA bonds (24-26), or by action on receptors which are directly sensitive to membrane signals, or through a wide range of other mechanisms such as mechanical stimulation of growth factors or their receptors, or both (27). To date, these consequences of the action of a constraint field have been described only qualitatively, without quantifying the effect, simply describing an increase vs decrease, or stimulation vs inhibition.

In addition, the mechanisms by which these biomechanical forces produce their effect, i.e. “*en bloc*”, have not been studied since the action of physical signals is extremely fast and not compartmentalized as signals are in a liquid medium (11).

The imposition of biomechanical stress enables a shift in study from a biological reductionist approach to one that examines the interaction between the constraint field and the cancer mass on a macroscopic level, tumor volume, and changes thereof.

The measurement of tumor volume as the target variable in this report has developed through two stages:

Firstly, a feasibility study was performed to assess the distribution in space of a mix of cells and ferric nanoparticles grafted subcutaneously.

Next, the design and implementation of the experiment to test Proof of Concept: The two groups subjected to a Magnetic Field Gradient (MFG) were placed between 2 identical magnets in repulsive mode (Fig. 1B).

The aim of the experimental system was to be able to apply a constraint field to modify the forces between the tumor and its environment. This effect was maximum to the left and minimum to the right due to the difference of distance between magnet and tumor.

If a difference in reduction in volume between left and right was found, it was then possible to draw two conclusions: that the mechanisms involved were similar, and that an estimate of the orders of magnitude of the acting forces could be obtained. It was then necessary to measure the two living surfaces of a tumor, excluding necrosis. The measurement was enabled by digitizing histological sections.

### Volume and surface measurements

Regular samples were monitored to quantitate the growth kinetics of overall volume and surfaces on histologic sections, including the viable tissue and the volume of central necrosis which is found in this type of subcutaneous tumor that has no or little angiogenic network.

The first measurement is performed *in vivo,* with a caliper, through the skin an converted in mm^3^ during the 21 treatment days from D 18 to D 30 and then up to 74 days when the last animals were sacrificed.

The second measurement is the *ex vivo* surface areas (Fig. 1C and 1D): left living tumor (located on the left in contact with the ipsilateral magnet and right living tumor located on the right and one diameter remote from magnet) and necrosis (all measured in mm^2^) on histological sections of tumors taken at different times of the experiment.

## MATERIALS AND METHODS

### Graft of MDA-MB-231 tumor in BALB/C Nude mice

Mice were 6-7 weeks old and weighed 16-20 g at reception (Charles River, Wilmington, MA). Ten million (1 x 10^7^) MDA-MB-231 tumor cells (American Type Culture Collection, Manassas, VA) were suspended in a volume of 0.3 mL RPMI 1640 medium containing either 5 mg of iron (300 μl of 100 nm nanoparticles/injection Fluid MAG D, Chemicell, Germany) or no iron nanoparticles. The appropriate suspensions were inoculated subcutaneously into the right and left flanks of 45 female athymic BALB/C *Nude* mice, irradiated 24-72 hours before with a γ-source (whole body irradiation, 2 Gy, ^60^Co, BioMEP Sarl, Bretenière, France) (28, 29).

### Animal husbandry and Ethics

Animals were living and personel was working in a Special Pathogen Free environment after irradiation. Viability and behavior were recorded every day. Body weight was recorded twice weekly.

Time of tumor cell injection was defined as Day 0 (D 0).

Animal experiments were performed according to English ethical guidelines and were approved by the Animal Care and Use Committee of Pharmacy and Medicine University of Dijon, France on 26 February 2012 (protocol N° 499). Treatment was performed under isoflurane anesthesia. Euthanasia was performed using cervical dislocation under isoflurane anesthesia.

All efforts were made to minimize suffering.

### Magnets

Magnet, cylindrical shape of 30 mm diameter and 100 mm height, maximum induction at magnet surface 0.66 Tesla (T) manufactured by Vakuum Schmeltz (Germany).

### Formation of groups and length of treatment

When tumors reached approximately 100-200 mm^3^ the mice were randomized into 4 groups after checking (ANOVA, p<0.05) that the mean volumes of the grafted tumors were similar.

Treatment involved a 2-hour exposure per day to a magnetic field gradient (MFG) for 21 days, from day 18 to day 39 for tumors with nanoparticles (Group 1; G1).

Group 2 (G2) had tumors with nanoparticles which were not exposed to the MFG;

Group 3 (G3) had tumors without nanoparticles, but were exposed to the MFG in the same way as G1;

Group 4 (G4) had tumors without nanoparticles which were not exposed to MFG;

G1 was defined as the treatment group;

G2, G3 and G4 are the control groups.

This is described in Table 1A.

**Table 1:**
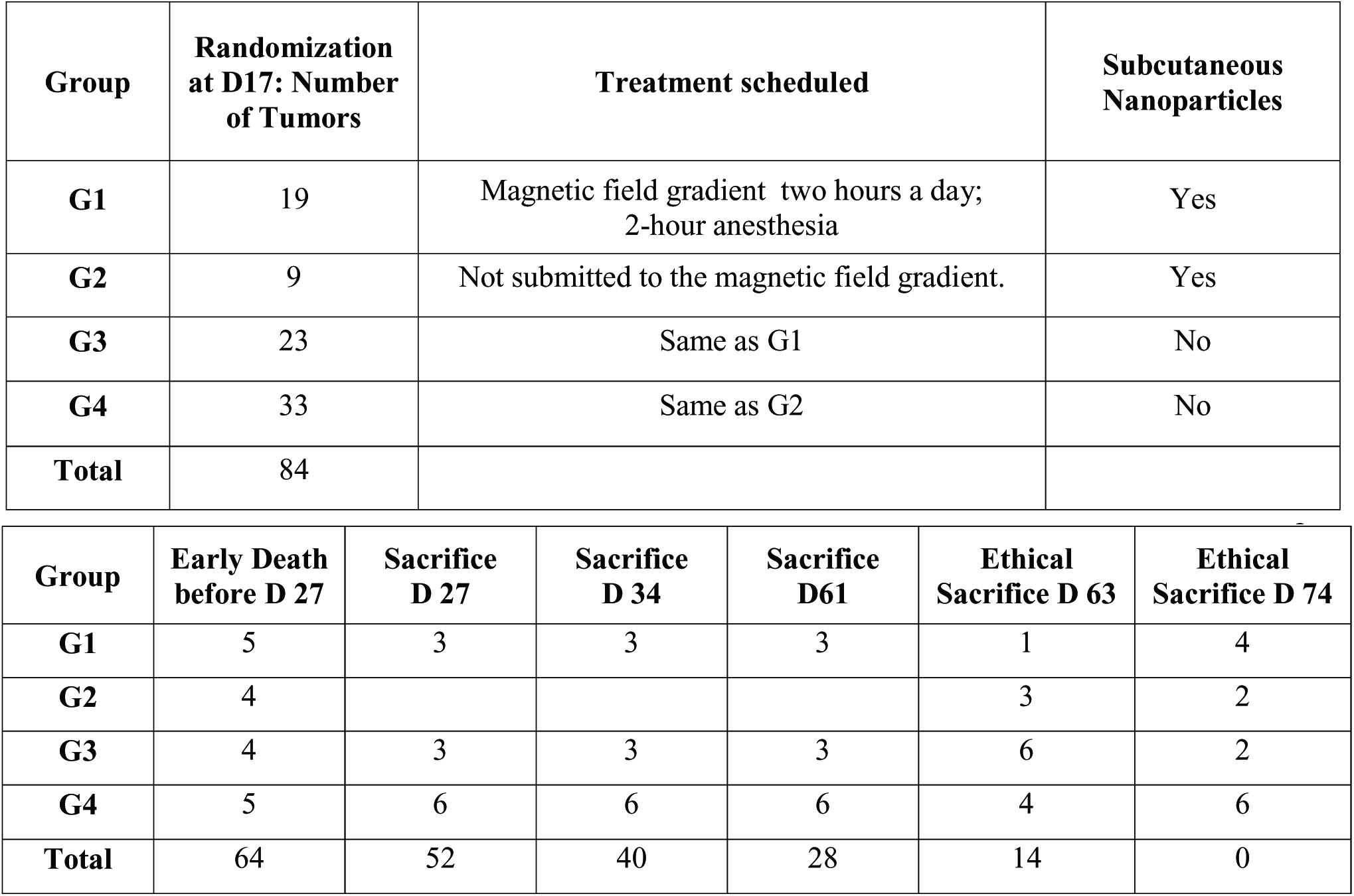
1A: Groups at randomization; Total number of tumors: 84 1B: Sampling of tumors during the experimentation

### Experimental Design

Samples of tumor were obtained by humanely killing the mice in order to monitor the change in tumor volumes over time by analyzing the maximum histological surface area of the tumors from which samples were taken.

Several samples were taken during the experiment: 3 tumor samples in G1 and G3 and 6 tumor samples in G4 on D 28, D34 and D 61.

It was not possible to obtain samples in G2 (no MFG, nanoparticles) which was made up of only 9 tumors.

The last sample was taken on D 61, i.e. 22 days after the end of treatment following a final *in vivo* caliper measurement on D 59.

The remaining living mice were humanely killed for ethical reasons on D 63 and D 74. The groups are described in Tables 1A and 1B.

**Table 2:**
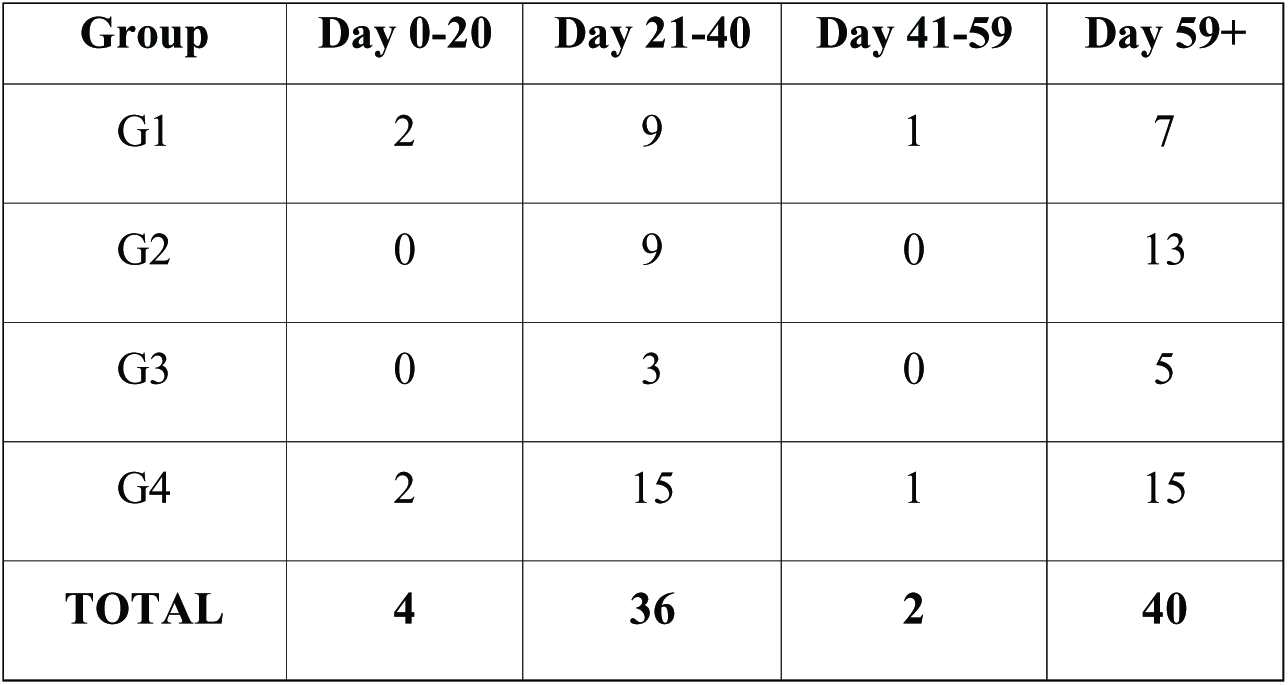
The distribution of tumor sampling every 10 days

The distribution of the 4 groups of tumors and change in populations over time are shown in Table 2.

### Estimate of force

A graph was available which displayed the magnetic field gradient in the airgap of the magnets (Altran, France). The force generated according to the amount of iron administered was approximately 10 Pa, 3 mm deep in the tumor, in contact with the magnet (on the left) and 0 to a few Pa on the right. This is illustrated in Fig 1E.

### Sample collection

During the course of the experiment, animals were humanely killed if any of the following signs occurred:

‐ Signs of suffering (cachexia, weakening),
‐ Compound toxicity (hunching, convulsions),
‐ Tumor ulcerating and remaining open,
‐ Position of the tumor interfering with movement/feeding,
‐ 15% body weight loss for 3 consecutive days or 20 % body weight loss for 1 day.

When the tumor volume reached a maximum volume of 2,000 mm^3^, an autopsy (macroscopic examination of heart, lungs, liver, spleen, kidneys and gastro-intestinal tract) was performed on all killed mice in the study, and if feasible, on all moribund mice of mice found dead. Autopsy observations were recorded.

All tumors were resected, fixed in formalin, and shipped to the pathologist.

Each paraffin section was cut at 4 μm, and the slides stained with HES, (hematoxylin, eosin and safranin).

### Treatment Randomization

Randomization before treatment was carried out on Day 17 and treatment began on Day 18. Treatment lasted 21 days. The animals subjected to an MFG were anaesthetized throughout their exposure to MFG, i.e. 2 hours per day, 7 days per week for 21 days. They were then observed.

### *In Vivo* Measurement

The length and width of the tumors were measured twice weekly in the 57 days of the experiment during the times when the tumors were easily measurable, i.e. D 17 to D 74.

At each measurement of tumor diameter, i.e. twice weekly, the airgap between the magnets was adjusted in order to maintain the left side of the tumor in contact with the left magnet and the right side of the tumor a distance of 1 diameter from the right magnet with a precision in the region of 0.1 mm.

The tumor volume (TV) was estimated by the formula: 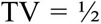 Length x Width^2^, expressed in mm^3^.

A first statistical analysis of the TV was performed on the tumors treated to D 39 and measured at least up to D 59 (last measurement before D 61 humane killing; D 59 population) and up to D 74 (last measurement before killing at D 63 and D 74; D 59+ population) (see Table 1B).

### *Ex vivo* Measurement, tumor surfaces measured on histological sections

A second statistical analysis was performed on the tumor surface measured either on the right or on the left side. Analysis was performed on “D 59” surfaces and on “D 59+”.

All of the tumors removed were digitized (Image J) on a Philips Ultra Scan 1.6 RA. The tumors were sectioned along their longest axis (see Fig. 2).

**Fig 2:**
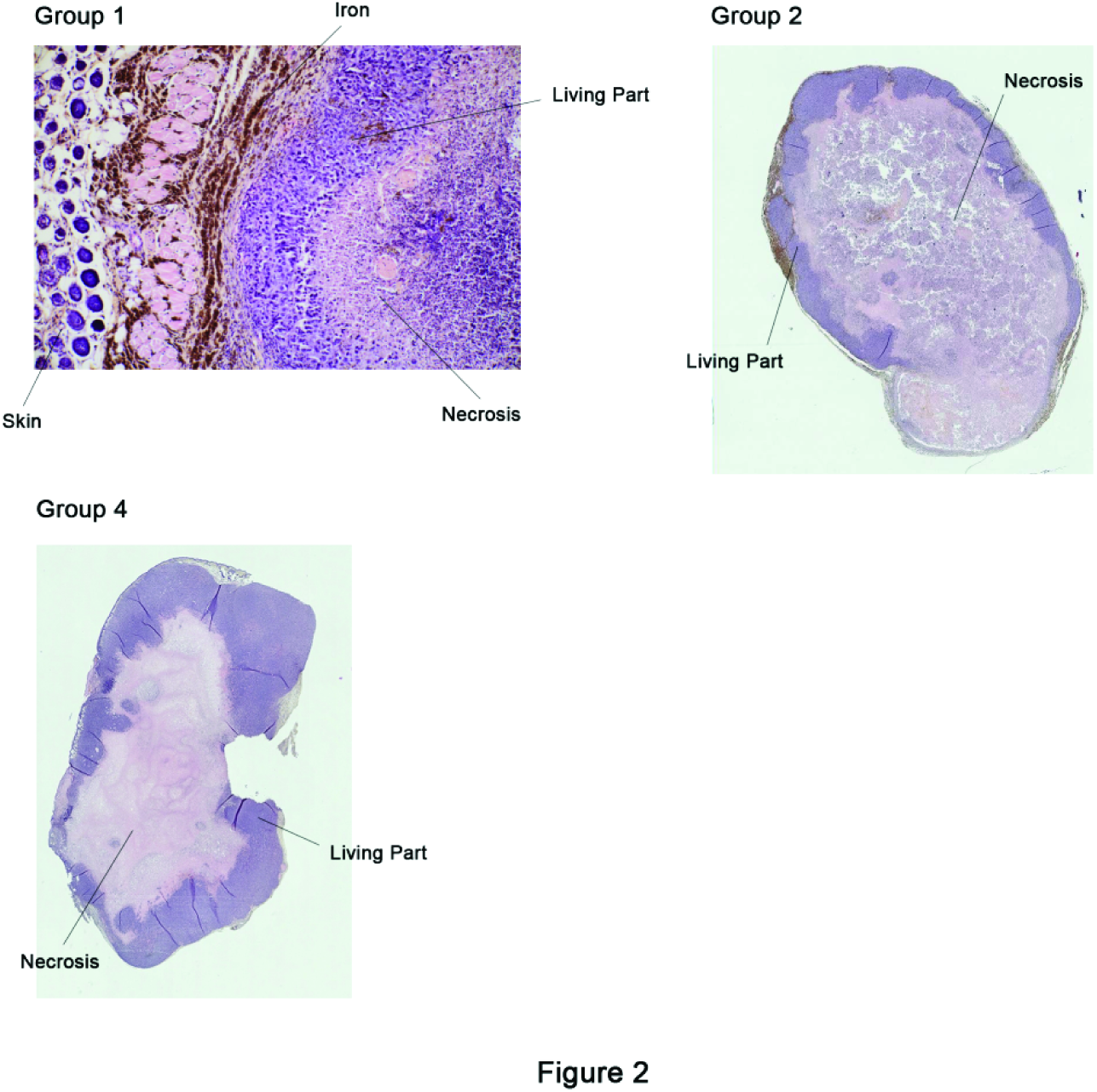
Fig. 2: sections of tumors. Staining: HES; Iron in black. Magnification: x50: Group 1 (treated); x5: Group 2 (nanoparticles, no MFG) and Group 4 (no MFG, no nanoparticles).

After digitization the tumors were divided into three parts:

Living part located on the right, living part located on the left and the necrotic part.

The delineation between the living parts was geometrical along the vertical of the long axis. The demarcation of necrosis was carried out under the supervision of the pathologist.

## STATISTICAL METHODS

All statistical analyses were performed using *in vivo* manager^®^ software (Biosystèmes, France). Statistical analyses of mean body weight changes, mean tumor volumes at randomization, mean target tumor volumes V, mean times to reach the target volume V and mean tumor doubling times were performed using the Bonferroni/Dunn test. All groups were compared with each other.

The following summary statistics were calculated for each variable (volume or surface area): number of tumors measured (n), range (minimum and maximum values), 25 %, 50 % (median) and 75 % quartiles and mean ± standard deviation.

Where no departure from the heteroscedasticity and normality assumptions was detected before or after log transformation of the data, a variable was statistically analyzed using a one-way ANOVA. The treated group was then compared to the mean of all 3 control groups by testing the following Helmert contrast: 1xG1-1/3x (G2+G3+G4). The rationale for merging control groups was the absence of statistically significant difference between the groups. Where the statistical assumptions of ANOVA were unmet, pairwise comparisons were performed using the Mann-Whitney U test. The same Helmert contrast was used as for the parametric procedure. Unless otherwise stated, a p value < 0.05 is considered as significant.

## RESULTS

The feasibility study showed that after the subcutaneous injection of a mix of nanoparticles (iron; 100 nm of diameter) and cells (MDA MB 231), nanoparticles distributed around the tumor, due to the large difference of free surface energy. This is illustrated in Fig. 3.

**Fig 3:**
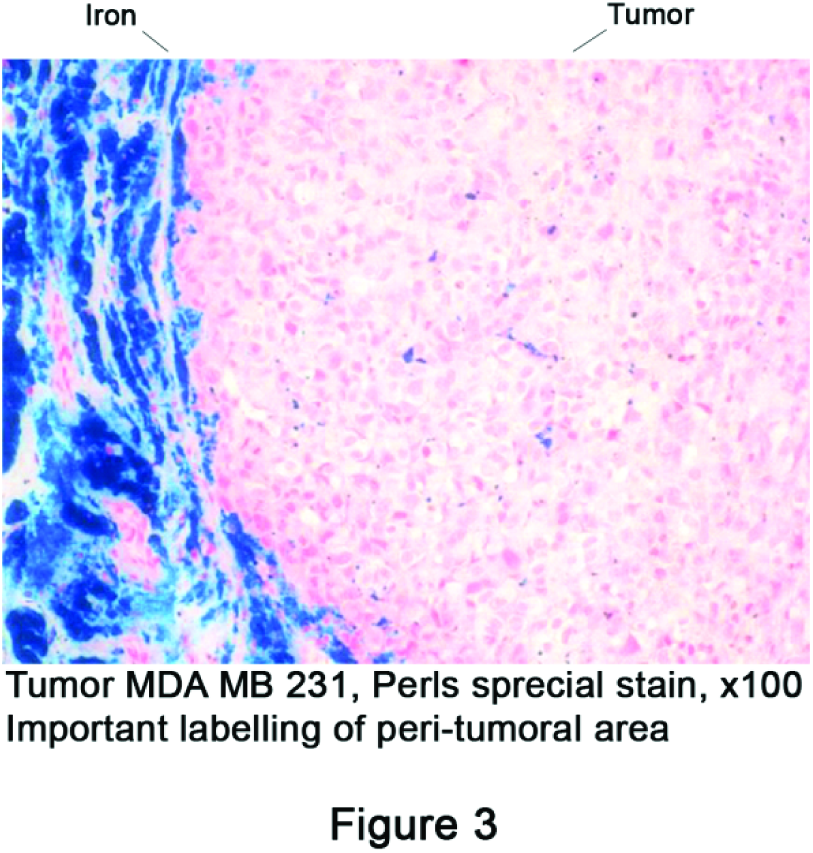
Fig. 3: Proof of Concept: physical feasibility *in vivo*. Spreading of the nanoparticles around a subcutaneous grafted tumor 21 days after injection of a nanoparticles/MDA MB 231 cell mix; Tumor in pink (HES), iron in black.

### Statistical results of the Proof of Concept

Seventy-nine out of the 82 tumors were included in the statistical analysis. The remaining 3 tumors could not be measured ^2^.

Forty-four sections were non-analyzable for technical reasons: This material is very fragile and shape was sometimes very unequal. There is no gross difference between the analyzable groups and the non-analyzable groups (see Table 3).

**Table 3:** The distribution of non-analyzable section by group

| Group | N | % |
| --- | --- | --- |
| G1 | 9/19 | 47 |
| G2 | 13/22 | 59 |
| G3 | 3/8 | 36 |
| G4 | 19/33 | 58 |

The results of volume and surface area measurements of measurable tumors are displayed in Table 4.

**Table 4:** Effect of a magnetic field gradient on *in vivo* tumor volume and *ex vivo* on its left and right surface areas

| Volumes are expressed in mm<sup>3</sup> and surface areas in mm<sup>2</sup> |  |  |  |  |  |
| --- | --- | --- | --- | --- | --- |
| Variable | Group | Range | Median (Q1;Q3) | Mean (SD) | Test used & p-value |
| **Volume,**<br>**D 59 tumors** | 1 (n=18) | 94; 966 | 246 (159; 529) | 383 (282) | Method 3, p=0.067<sub>NS</sub> |
|  | 2 (n=21) | 96; 2106 | 1334 (334; 1784) | 1086 (756) | a |
|  | 3 (n=8) | 87; 1089 | 596 (161; 914) | 565 (395) | a |
|  | 4 (n=32) | 70; 1996 | 388 (175; 928) | 635 (613) | a |
| **Volume,**<br>**D 59+ tumors** | 1 (n=7) | 346; 966 | 529 (502; 840) | 647 | Method 3, p=0.015 |
|  | 2,3,4 (n=33) | 256; 2106 | 1334 (758; 1784) | 1250 | - |
| **Surface,**<br>**D 59 tumors,**<br>**Left side** | 1 (n=9) | 3.0; 42.4 | 5.6 (4.2; 9.8) | 10.9 (12.8) | Method 2, p=0.005<sub>**</sub> |
|  | 2 (n=14) | 4.9; 34.3 | 20.8 (16.3; 26.4) | 20.6 (8.7) | b |
|  | 3 (n=4) | 13.2; 23.2 | 16.5 (13.6; 21.0) | 17.3 (4.6) | b |
|  | 4 (n=19) | 1.8; 55.5 | 14.8 (9.0; 21.0) | 17.1 (12.5) | b |
| **Surface,**<br>**D59+ tumors,**<br>**Left side** | 1 (n=4) | 5.6; 42.4 | 14.2 (7.7; 30.5) | 19.1 (16.5) | Method 2, p=0.172<sub>NS</sub> |
|  | 2 (n=10) | 16.3; 34.3 | 23.8 (19.7; 28.8) | 24.6 (6.1) | c |
|  | 3 (n=4) | 13.2; 23.2 | 16.5 (13.6; 21.0) | 17.3 (4.6) | c |
|  | 4 (n=10) | 14.8; 55.5 | 20.5 (18.1; 27.3) | 25.1 (12.2) | c |
| **Surface,**<br>**D 59 tumors,**<br>**Right side** | 1 (n=9) | 1.9; 29.2 | 4.7 (3.6; 7.2) | 7.7 (8.4) | Method 2, p=0.001<sub>**</sub> |
|  | 2 (n=14) | 5.2; 55.0 | 25.0 (11.7; 28.4) | 23.1 (14.0) | d |
|  | 3 (n=4) | 14.6; 35.2 | 18.0 (15.5; 27.3) | 21.4 (9.4) | d |
|  | 4 (n=19) | 1.5; 51.8 | 12.5 (6.0; 28.6) | 18.7 (14.9) | d |
| **Surface,**<br>**D 59+ tumors,**<br>**Right side** | 1 (n=4) | 4.6; 29.2 | 8.6 (5.9; 19.6) | 12.8 (11.2) | Method 1, p=0.093<sub>NS</sub> |
|  | 2 (n=10) | 11.8; 55.0 | 26.1 (24.8; 33.5) | 29.1 (11.9) | e |
|  | 3 (n=4) | 14.6; 35.2 | 18.0 (15.5; 27.3) | 21.4 (9.4) | e |
|  | 4 (n=10) | 10.9; 51.8 | 28.1 (12.5; 40.3) | 28.6 (14.0) | e |

#### Method 1

One way ANOVA on untransformed data. In case of significance, the treated group was compared to the mean of all 3 control groups by testing the following Helmert contrast: 1xG1-1/3x (G2+G3+G4).

#### Method 2

Same as method 1 but using log-transformed data.

#### Method 3

The treated group was compared to the mean of all 3 control groups using the Mann-Whitney U by test.

##### Q1; Q3

First and third quartiles.

##### a,b,c,d,e

Mean values with the same letter were similar. No statistically significant differences were detected between the control groups using pair wise comparisons with Bonferri-Holm adjustment.

### *In vivo* volume

As shown on Fig. 4, median volume on D 59+ was significantly lower in G1 (529 [346; 966]) mm^3^, p=0.015) than in G2, G3 and G4 (1334 [256; 2106]).

**Fig 4:**
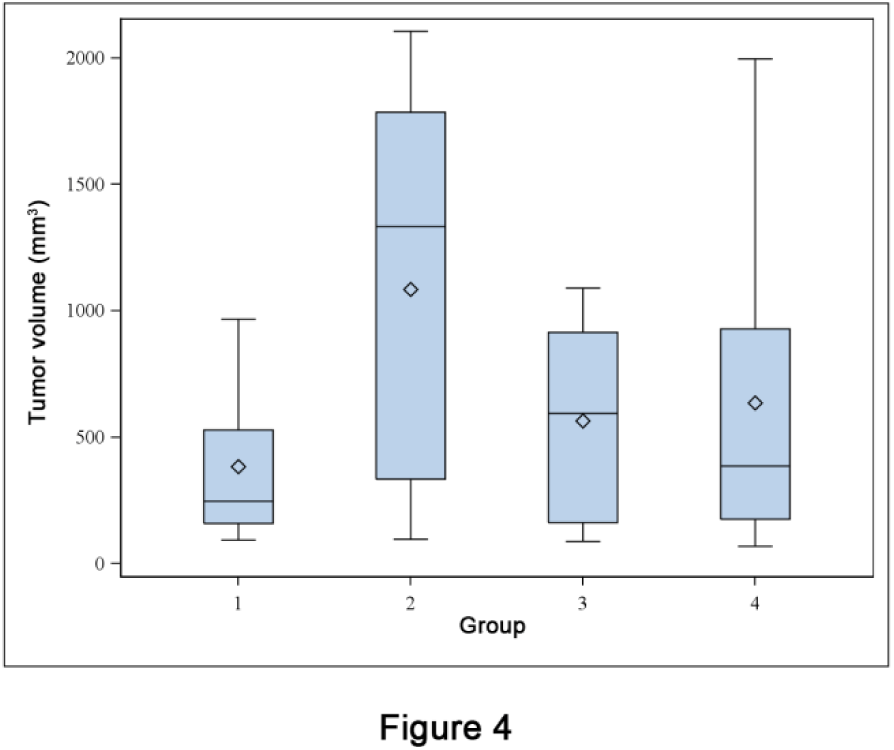
Fig. 4: D 59+ tumors. Boxplot of tumor volume (mm^3^) in the treatment group (1) and the three control groups (2 to 4). The bottom and top of the blue box are the first and third quartiles, and the horizontal rule inside the box is the second quartile (median). Whiskers correspond to ± 1.5 IQR (interquartile range). Means are plotted with an open diamond.

The tumor growth curve for the D 59+ population *in vivo* is shown in Fig. 5.

**Fig 5:**
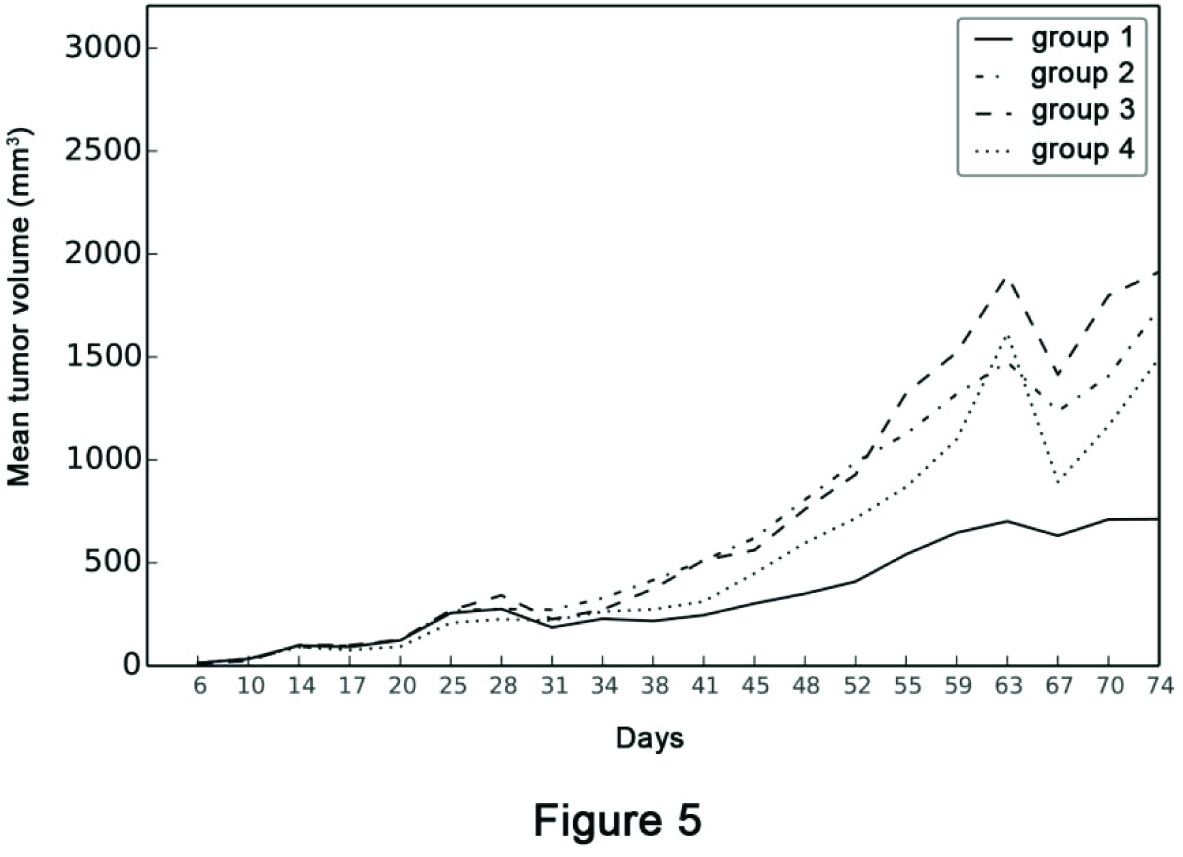
Fig. 5: Evolution of the growth of tumors between D 0 (graft) to D74 (last sacrifices). Divergence begins on D 31 at the end of the treatment.

Median [followed by range] volume of D 59 tumors was 246 [94; 966], 1334 [96; 2106], 596 [87; 1089], and 388 [70; 1996] mm^3^ in groups G1, G2, G3 and G4 respectively. No statistically significant difference was detected between the groups (p=0.067).

### *Ex vivo* surfaces

As shown on Fig. 6A, median left surface of all tumors was significantly lower in G1 (5.6 [3.0; 42.4] mm^2^, p=0.005) than in G2 (20.8 [4.9; 34.3]), G3 (16.5 [13.2; 23.2]), and G4 (14.8 [1.8, 55.5]).

**Fig 6:**
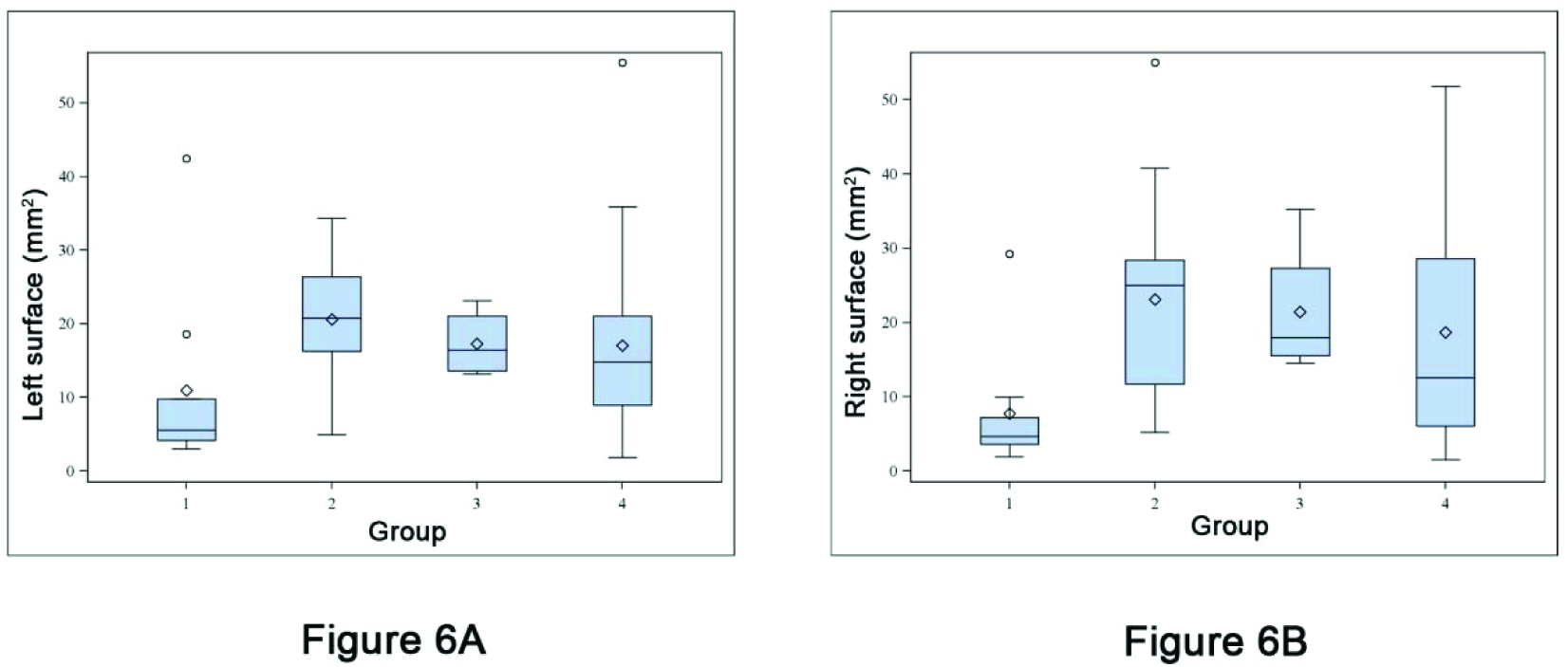
Fig. 6A and 6B: Comparison between surfaces. 6A: Bloxpot of left tumor surface in treatment group (1) and the three control groups (2 to 4). The bottom and top of the blue box are the first and third quartiles and the horizontal rule inside the box is the second quartile (median). Whiskers correspond to ± 1.5 IQR (interquartile range). Data not included between the whiskers are plotted as outliers with an open circle. Means are plotted with an open diamond. 6B: Boxplot of right surface.

Median left surface on D 59+ was 14.2 [5.6; 42.4], 25.0 [5.2; 55.0], 18.0 [14.6; 35.2], and 12.5 [1.5, 51.8] mm^2^ in groups G1, G2, G3 and G4 respectively. No statistically significant difference was detected between the groups (p=0.172).

As shown on Fig. 6B, median right surface of all tumors was significantly lower in G1 (4.7 [1.9; 29.2] mm^2^, p=0.005) than in G2 (25.0 [5.2; 55.0]), G3 (18.0 [14.6; 35.2]), and G4 (12.5 [1.5, 51.8]).

Median right surface on D59+ was 8.6 [4.6; 29.2], 26.1 [11.8; 55.0], 18.0 [14.6; 35.2], and 28.1 [10.9, 51.8] mm^2^ in groups G1, G2, G3 and G4 respectively. No statistically significant difference was detected between the groups (p=0.093).

### Toxicity

Some deaths occurred. Twenty (20) mice died after treatment administration or were found dead: 6 in G1, 4 in G2, 4 in G3 and 6 in G4. They were not replaced.

In G1 four mice were found dead after treatment administration (at D 19, 20, 22 and 41) and two mice were found dead in their cage (at D 21 and 22).

In G3 two mice were found dead after treatment administration (at D 18 and 22) and two mice were found dead in their cage (at D 33 and 45).

In G2, 4 mice were found dead in their cage (at D 18, 21, 22 and 25).

In G4, 6 mice were found dead in their cage (at D 17, 18, 18, 20, 22, 41).

### Autopsy

All thoracic and abdominal organs were analyzed. The reports mentioned “Death probably due to Isoflurane” (when appropriate) or “No evidence of toxicity or suffering due to the tumor or weight loss”. No iron was seen in the macrophages of the autopsied organs.

## DISCUSSION

The aim of this study was to provide *in vivo* experimental evidence of the possibility of influencing tumor growth by biomechanical signals. This was based on results obtained *in vitro* in which the effect has been extensively demonstrated.

Two analyses were performed, one on the *in vivo* measurements of the volume of tumors grafted subcutaneously and the other on *ex vivo* digitized surfaces of tumors sampled.

A highly significant difference (p=0.015) was found in tumor volume measured *in vivo* in the D 59 + population of tumors after 21 days of treatment. The difference did not reach statistical significance (p=0.067) for all tumors included.

The analysis of the maximal histological surface comparing either the left side of tumors where the field of constraint was maximal or the right side where the field of constraint was minimal showed a significant difference between treated tumors compared to controls for all the D 59 population. No significant difference between surfaces right or left could be detected for D 59+ population.

The forces and pressures generated by the constraint field applied to the tumor produced complex effects related to the physical properties of the different constituent media, epithelial tissue, fibrosis, resistance of part of the ECM in contact with the tumor tissue and more remote tissues. It has been shown recently (30) that pressure exerted by a growing tumor influences biologically and mechanically the normal tissue at a distance.

These effects result from mechanics which have hardly been explored at all to date. This study does not answer the question of describing or measuring traction, impulses, shear and other forces involved.

The right/left comparison, however, may give a clue to answer the question of biomechanical impact resulting in highly significant surface differences between the left, the region exposed to the greatest pressure, and the right, the region of the lowest pressure. An increase in pressures applied would therefore appear to have a greater effect on tumor growth, despite the low level of forces exerted on tumor.

### The future

After validating this Proof of Concept for this experimental device which cannot be used in a clinical situation. Moreover, the lesser increase of tumor volume is a low marker of anticancer activity. Proof of Efficacy using more powerful devices is a logical next step.

There are four available parameters than can be varied: magnetic field of the magnet, distance to the tumor, quantity of iron and approximate stiffness as derived from literature. There is, therefore the basis for further mechanistic studies.

The experimental design will be based on iron deposition in neoangiogenesis. To do this, specific neoangiogenesis nanoparticles already developed and used in animal imaging will be employed. The geometry of the iron deposit around the tumor makes it possible to apply a constraint field localized in the tumor with a turning ant tunable magnetic field gradient generated by electromagnets. Pressures could therefore be increased by a large factor.

Two problems arise: the first is constructing an experimental Proof of Efficacy model which may contribute information about the physical mechanisms involved on a macroscopic scale applicable to the cancer and its two tissues: Tumor and ECM, and to the surrounding normal tissue. This current model will allow insight to improve and better understand the experimental model chosen for Proof of Efficacy, i.e. orthotopic grafting of human pancreatic cancer into the pancreas of mice or human liver cancer into the liver of mice allowing an extrapolation to be made to the clinical situations of these locally advanced tumors which have no satisfactory treatment.

The second problem which arises is the microscopic analysis of biochemical signals generated by the physical changes. Description of signal transduction pathways and the expression of common promoter and inhibitor gene products under the influence of biomechanical forces would be of both academic and “product development” interest. Correlation with the observations of tumor growth as described herein could provide a model for targeted intervention designed to mimic the biomechanical effects.

Mechanical signals are transmitted from the ECM to the nucleus through CSK (31-33). The CSK components, actin filaments, microtubules and microfilaments, associated with mitochondria, representing approximately 20% of the cell volume, are semi-solid and in continuity with the tissue environment. They maintain architectural integrity of tissue (34) and mitochondrial thermodynamic function related to the CSK (35).

They are the support for the cell architecture through a reciprocal mechanism of compression and tension between them and the ECM or Basal Membrane (BM) (36, 37). This semi-solid system is changed greatly by mechanical cues during the process or carcinogenesis and is the most accessible part to analysis and change by physical means. Resonance frequencies of the CSK will be looked for.

## CONCLUSION

The study of tumor transformation through the action of forces and pressures and the introduction of the notion of constraint field in tissue is quite recent in oncology and also is relevant to embryogenesis (38), morphogenesis (39, 40), and tissue engineering.

This biomechanical framework makes it possible to redefine cancer as a disease of the cellular and, particularly, the tissue architecture leading to changes in cell metabolism ranging from oxidation to glycolysis and, ultimately, to a profound change in the genetic machinery (41).

This is valid for the 3 steps of carcinogenesis, from BM crossing to metastatic invasion by adeno- or squamous carcinomas, in other words a very large majority of cancers of epithelial origin.

The two medical devices used in this Proof of Efficacy, ferric nanoparticles (developed as MRI IV contrast media of neovessels) and magnetic field gradient generators are commonly used in rodents and patients without significant toxicity. Moreover until now physical oncology has been used as a research and diagnostic tool and its therapeutic use has only been imagined (42). This disruptive innovation was made possible by the *in vivo* construction of geometrical spreading of nanoparticles around the tumor. The same conditions will be used for a future Proof of Efficacy study before potential application to locally advanced epithelial tumors.

We can therefore conceive of producing a local well tolerated treatment centered on restoring normal biological properties to cancer tissue using physical manipulation.

## ACKNOWLEDGMENTS

The authors would like to thank Ms Emmanuelle Fourme for her help in statistics, M. Bertrand Cahuzac for his help in Physics, Ms Eléonore Lamoglia for the illustrations, Drs Ann Barrett (UK) and Roy Weiner (US) for their kind proofreading and Christelle Boileau for her assistance in the manuscript preparation.

## Footnotes

1 A patent application has been filed (PCT EP 2014 064995, Cell Constraint … Cancer Inc., France) to protect the applications of these findings.

2 The material is very fragile and was damaged during transportation

